## Appendix S1 for "Positive and negative effects of land abandonment on butterfly communities revealed by a hierarchical sampling design across climatic regions"

Appendix S1 Details in predictive model of species-specific responses to land abandonment

A hierarchical modelling approach is a powerful tool for predicting species-specific responses to environmental changes (Öckinger et al., 2010), and allows us to evaluate the broad-scale distribution of a species. Here, we developed a predictive model based on the habitat preferences of a species. As most of the species inhabit several habitat types, the predictive model was a multivariate extension of Eqn. 2:
*μ_β_*_1_*_i_* = *α*_0_ + *Σ_k_α_k_H_ik_*.	Eqn. S1
However, models that are too complex are vulnerable to overfitting and tend to have large prediction error. In such a situation, ridge regularization is useful to constrain overfitting and improve prediction accuracy (Hooten and Hobbs, 2015). In a Bayesian manner, ridge regularization can be simply implemented using the same normal prior, N(0, *σ_α_*^2^), for each regression coefficient. This prior works as a penalty for the divergence of squared regression coefficients from zero, and the standard deviation, *σ_α_*, corresponds to the weight of the penalty. The prior of *σ_α_* was set to a weakly informative prior, half-Cauchy(0, 5), and *σ_α_* was determined by calculating the posterior distribution. The prior of *α*_0_ was a normal distribution with mean 0 and variance 100. Samples from the posterior distribution were obtained using RStan 2.21.1 (Stan Development Team, 2016), with the same MCMC sampling settings and posterior diagnostics as in Sections (1) and (2).

To make projection maps for the effect of land abandonment on butterfly assemblages, we defined the loss (CL) and gain (CG) in butterfly species due to land abandonment as follows:

$$\mathrm{CL}_{j} = \sum_{i}^{N_{sp}} \delta_{ij}\left| min\left( \mu_{\beta1i},0 \right) \right|,$$

and

$$\mathrm{CG}_{j} = \sum_{i}^{N_{sp}} \delta_{ij}max\left( \mu_{\beta1i},0 \right),$$

where *N_sp_* is the number of species and *δ_ij_* is a binary indicator of whether the *i*th species is present in the *j*th grid cell. Operators $min\left( \mu_{\beta1i},0 \right)$ and $max\left( \mu_{\beta1i},0 \right)$ are filters to identify species affected negatively and positively, respectively. CL and CG can be interpreted as log-odds ratios of occurrence probabilities across species after land abandonment. Large values of CL or CG respectively indicate a large loss or gain in species in the local butterfly assemblage due to land abandonment.

We used nation-wide range maps of 70 butterfly species in Japan (Table S3), which were projected by Kasada et al. (Kasada et al., 2017) using distribution models (MaxEnt) from butterfly records obtained from the fourth and fifth National Surveys on the Natural Environment. Before calculating CL and CG, we excluded seven alpine butterflies, *Carterocephalus palaemon*, *Erebia neriene*, *E. ligea*, *Oeneis norna*, *Colias palaeno*, *Anthocharis cardamines* and *Aporia hippia*, because land abandonment is not relevant in their ranges. The range maps comprise presence and absence data for species in each grid cell of a “1 × 1 km mesh (the third mesh),” a Japanese national grid system whose unit cell size is 30′′ in latitude and 45′′ in longitude, approximately 1 × 1 km. We determined the expected coefficient of land abandonment, $\mu_{\beta1}$, using Eqn. S1, for 70 species. We then evaluated CL and CG separately for all species and for species classified as “near threatened (NT)”, “vulnerable (VU)”, “endangered (EN)” or “critically endangered (CR)” in the Red List of Japan in 2019 (Ministry of Environment, 2019). Then, we calculated CG – CL to evaluate whether loss of species richness due to land abandonment outweighed the gain.

Stan Development Team, 2016. RStan: the R interface to Stan. R package version 2.14.1.
