## Supplementary material for "Positive and negative effects of land abandonment on butterfly communities revealed by a hierarchical sampling design across climatic regions": Fig. S1

Fig. S1 Predicted effect of abandonment (posterior mean and 95% CI) using ridge regression. Blue dots and error bars indicate species whose upper 95% CI was below 0.

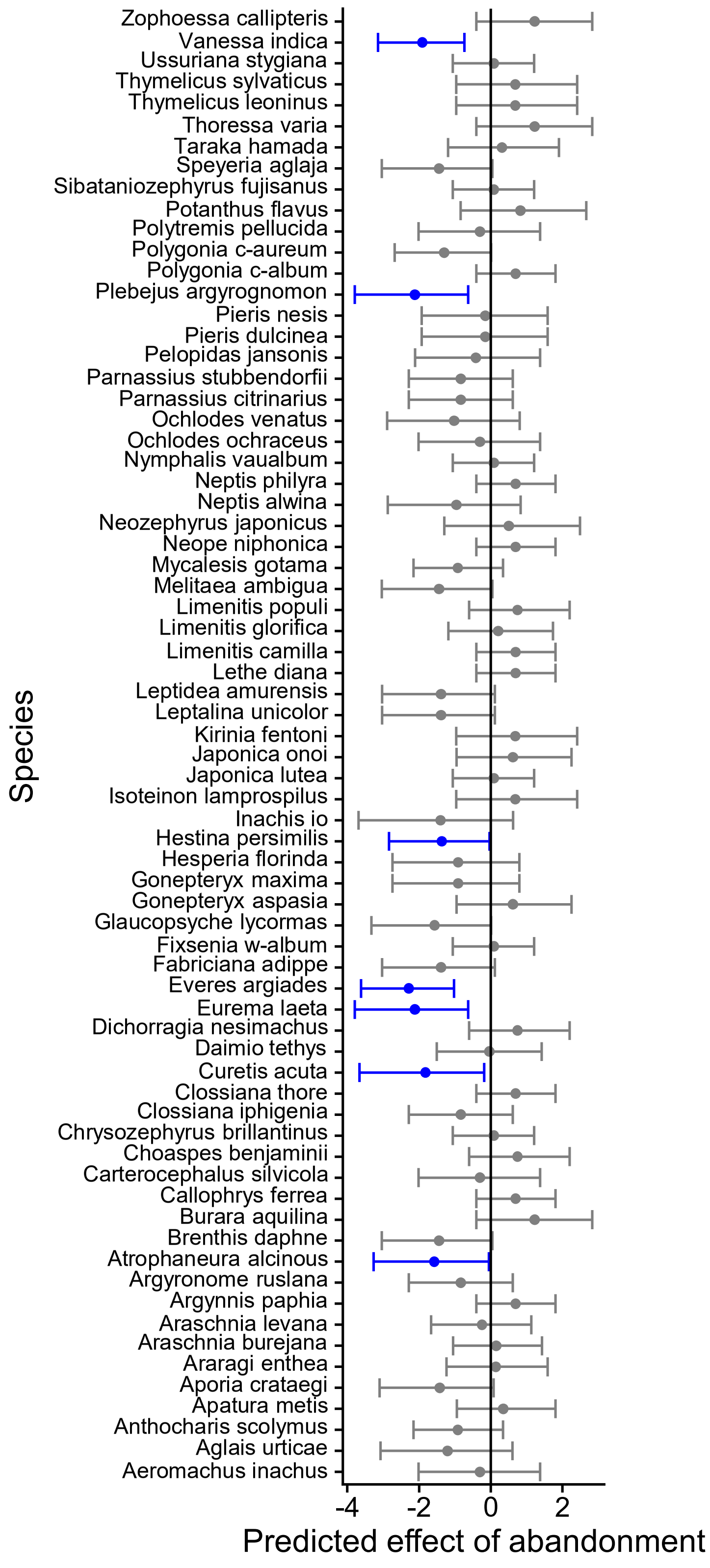

Fig. S2 Standard deviation of cumulative loss (CL, a), cumulative gain (CG, b), the proportions of CL (c) and CG (d) and the difference between CG and CL (CG - CL, e) in butterfly species richness due to land abandonment for all species.

(a)

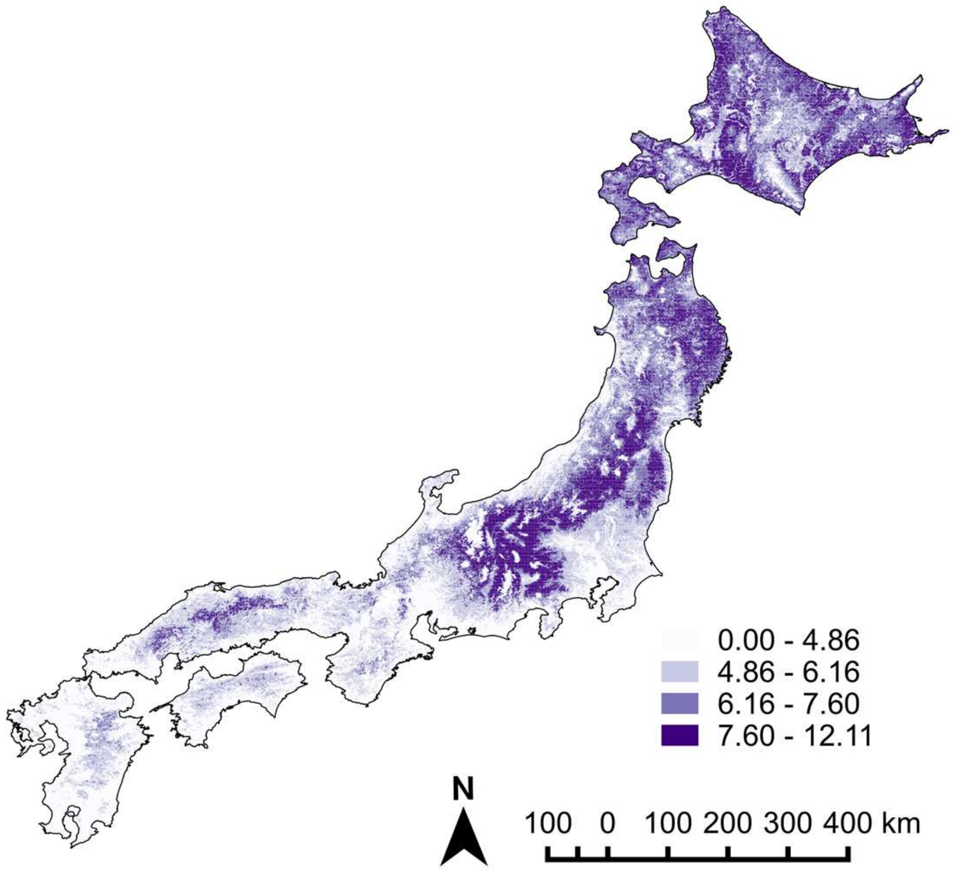


(b)

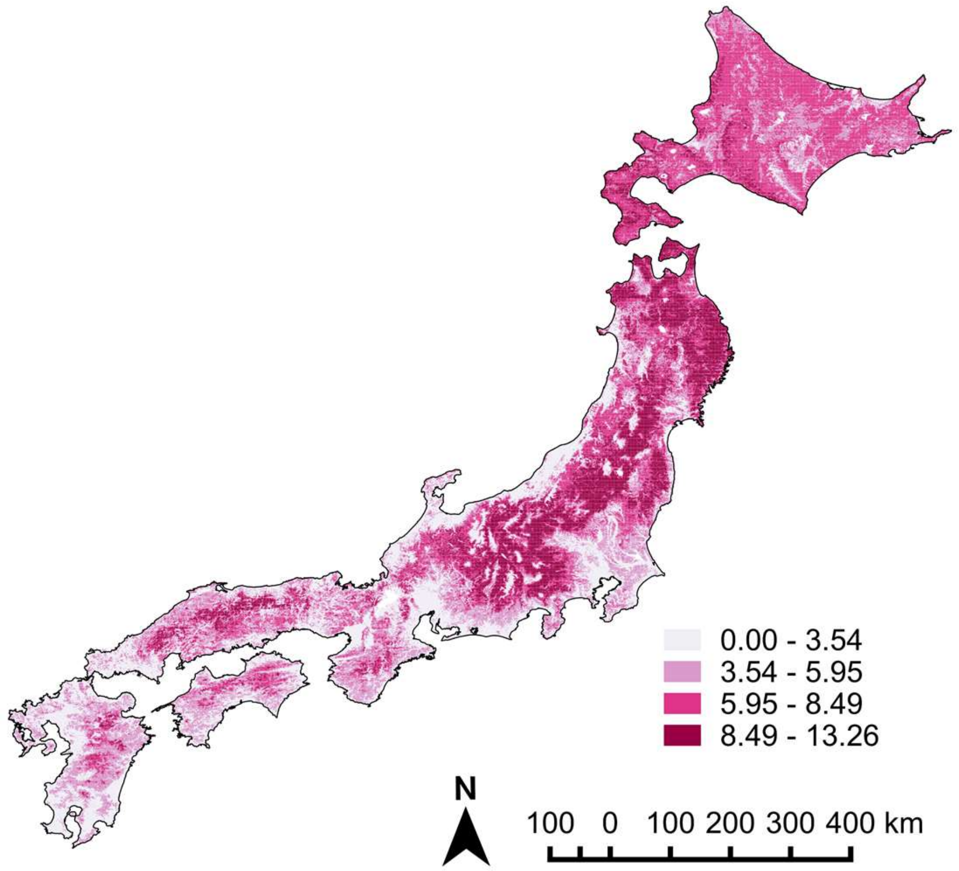

Fig. S2 continued.

(c)

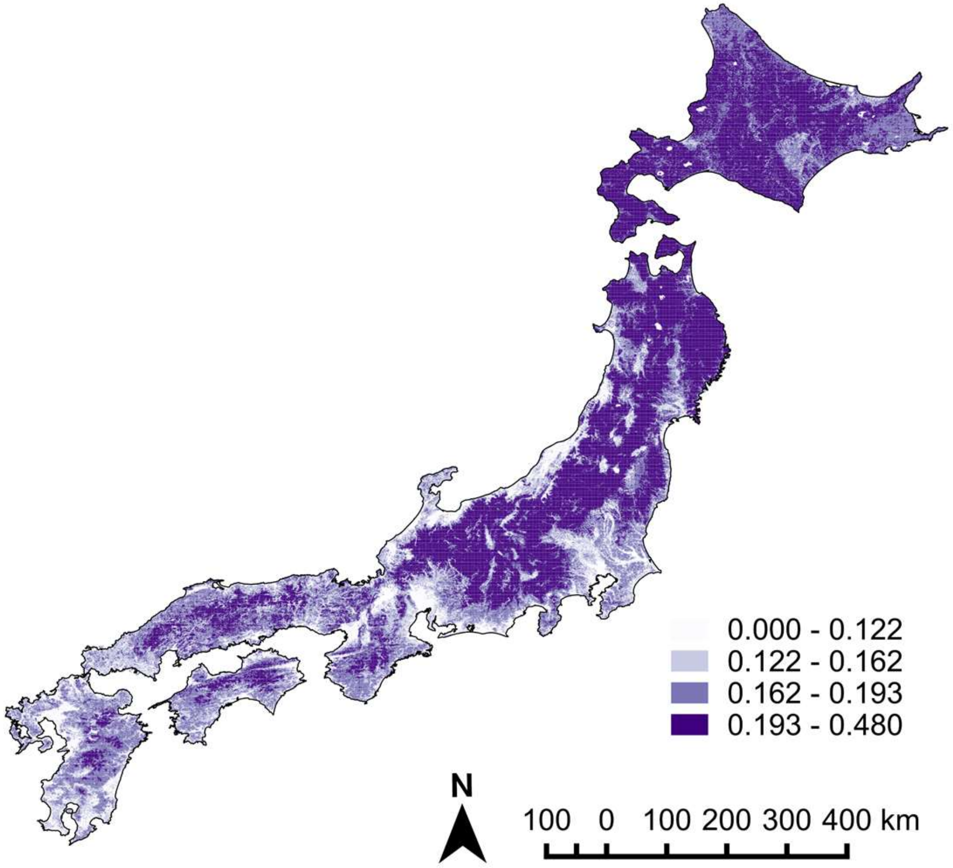


(d)

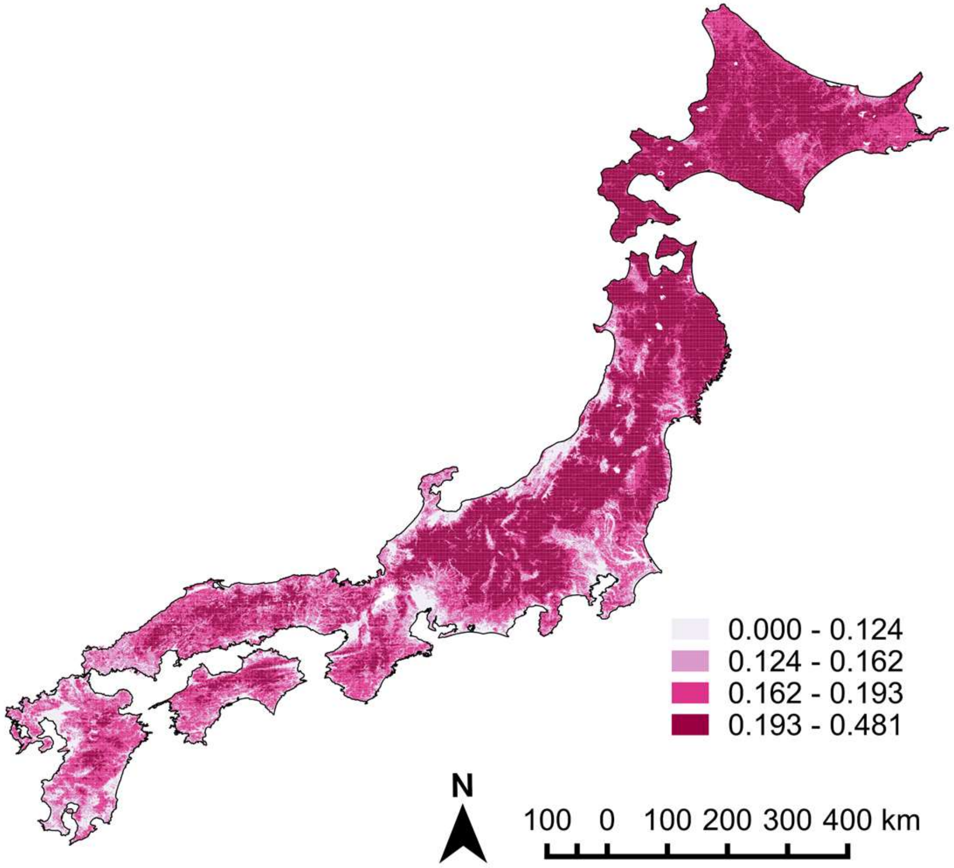

Fig. S2 continued.

(e)

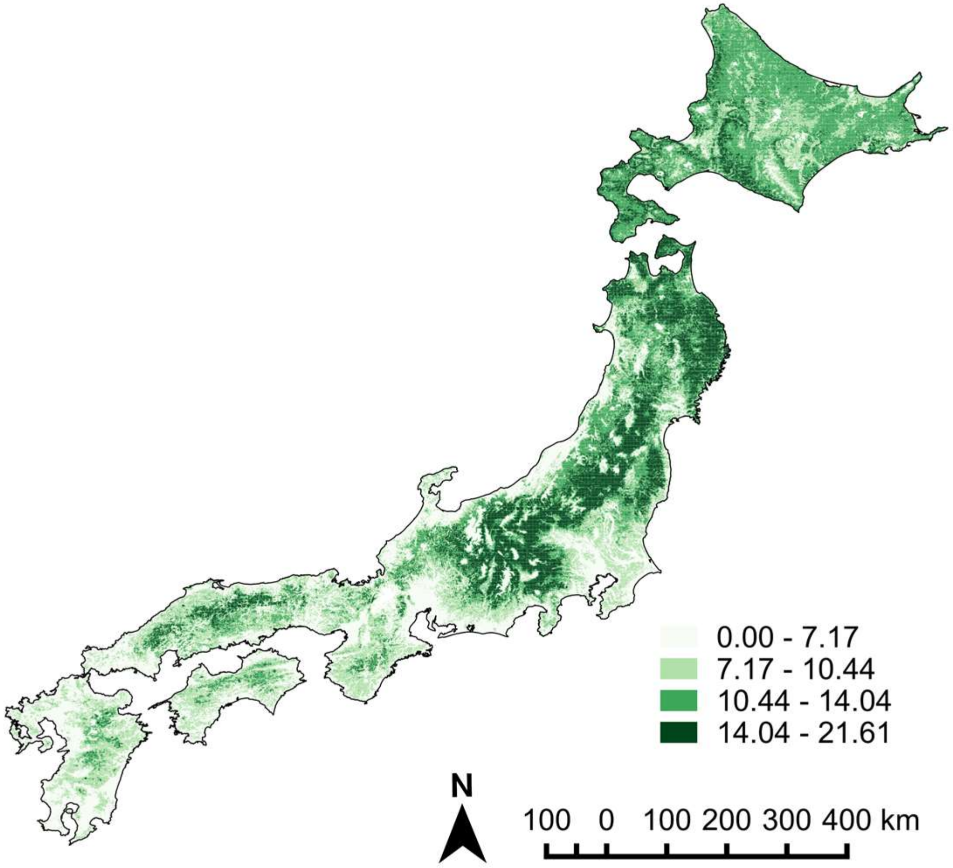

Fig. S3 Maps of cumulative loss (CL, a), cumulative gain (CG, b), the proportions of CL (c) and CG (d) and the difference between CG and CL (CG - CL, e) in butterfly species richness due to land abandonment for Red List species. The boxed graphs are hexagonal binning plots showing their relationship with elevation.

(a)

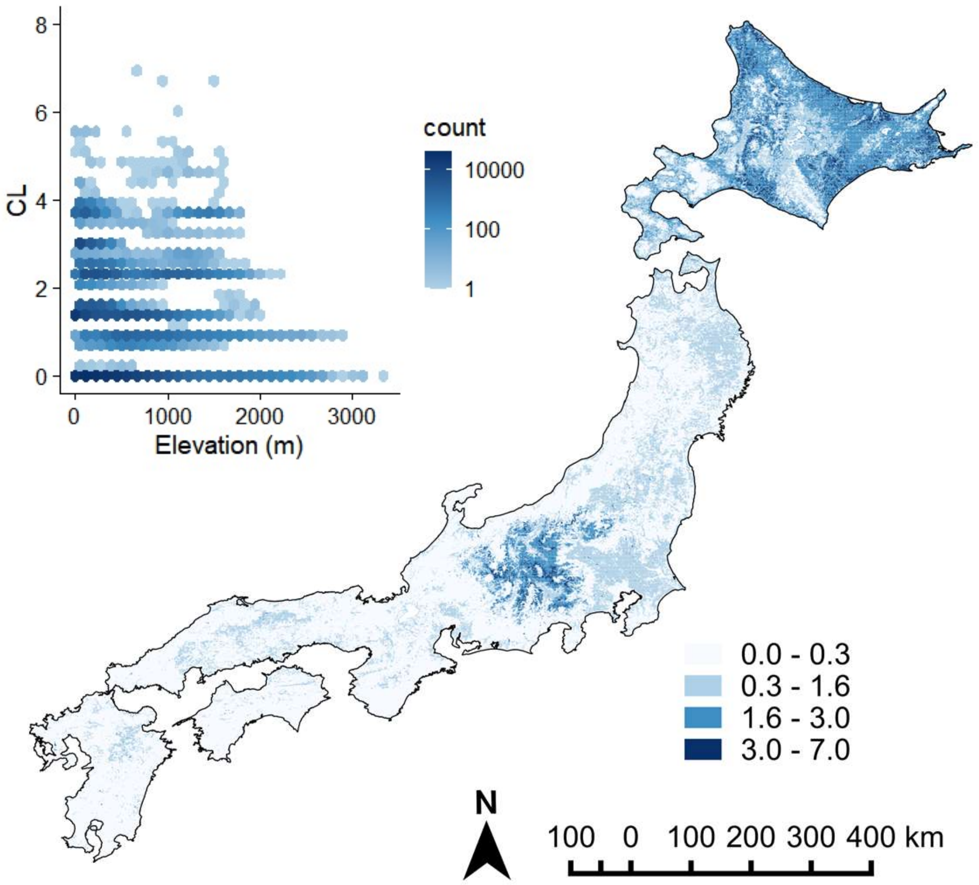


(b)

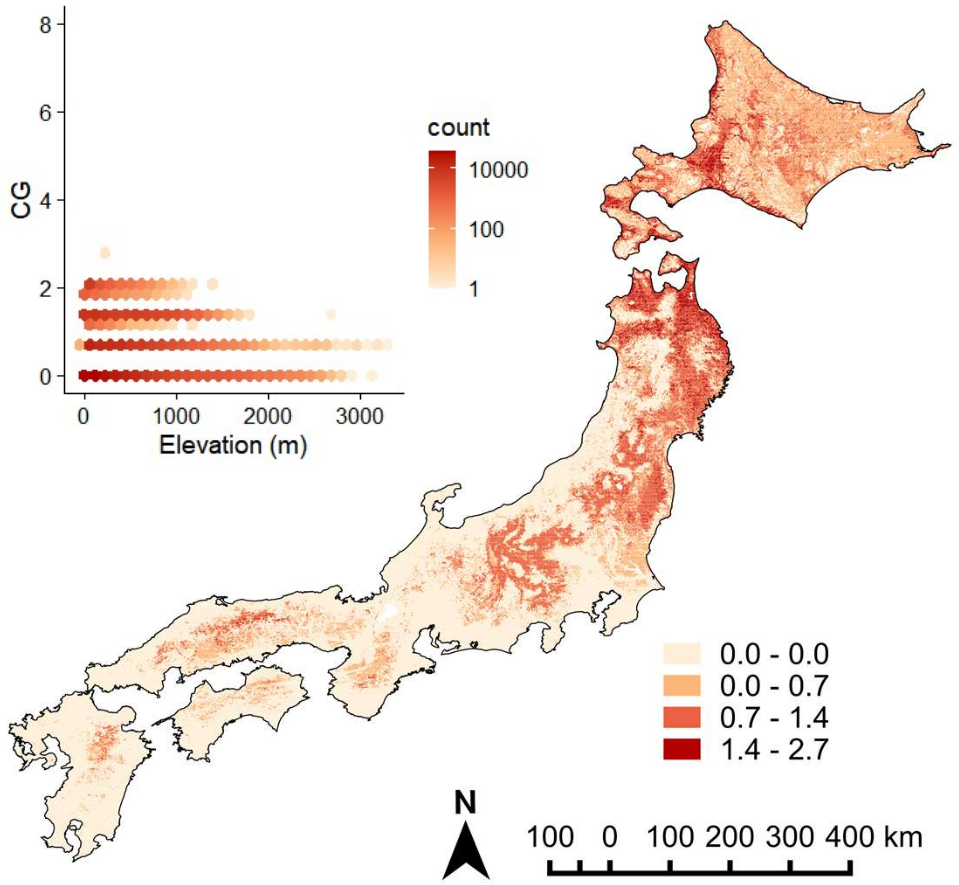

Fig. S3 *Continued.*

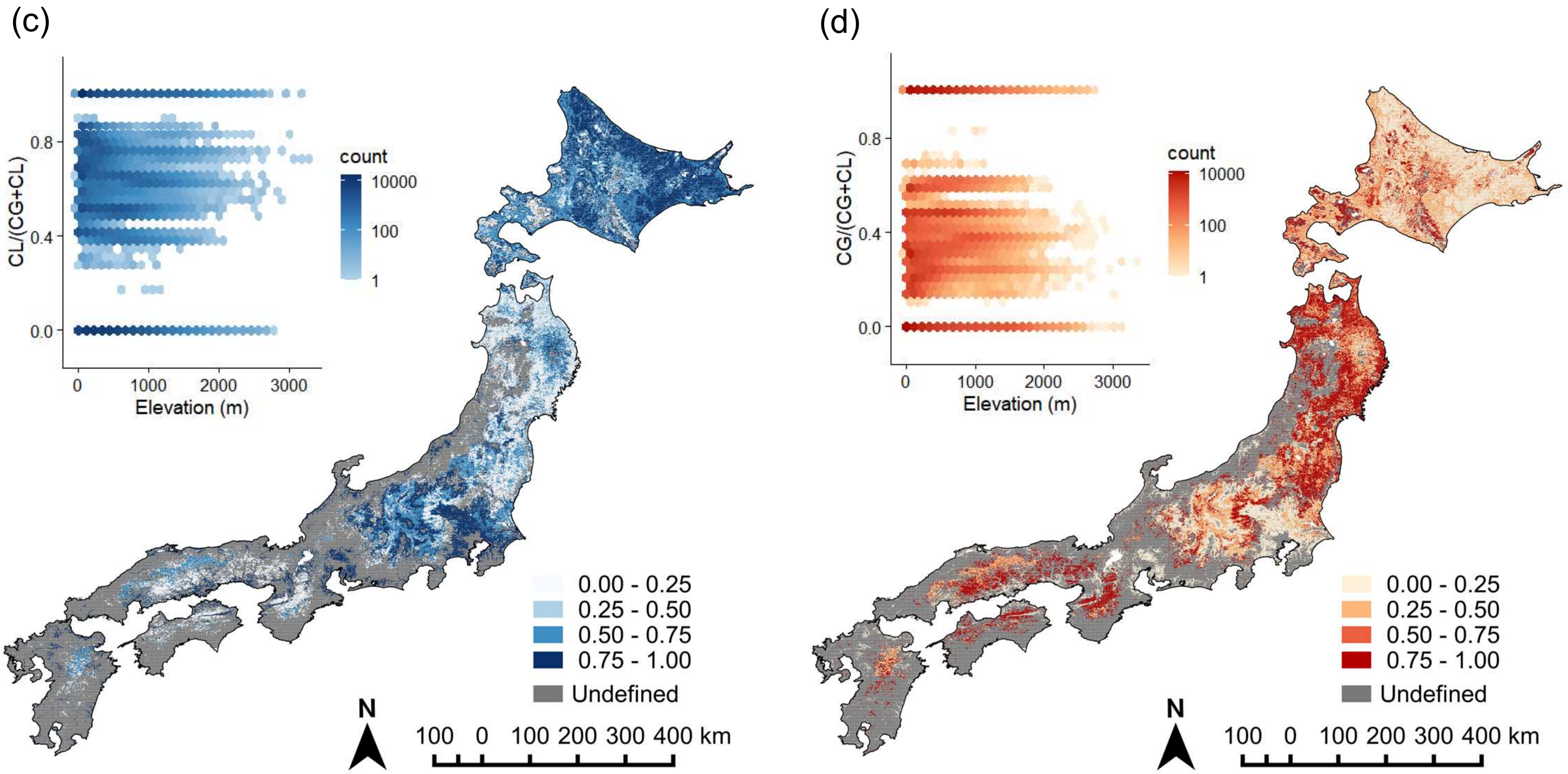

Fig. S3 *Continued.*

(e)

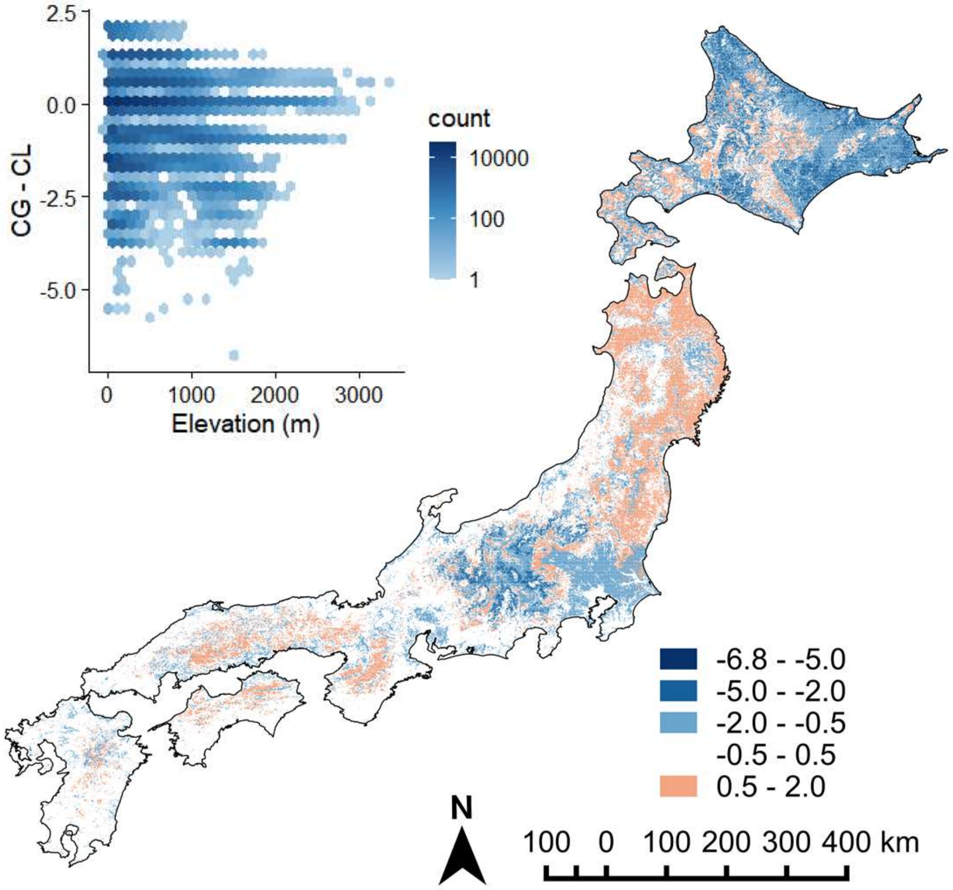

Fig. S4 Standard deviation of cumulative loss (CL, a), cumulative gain (CG, b), the proportions of CL (c) and CG (d) and the difference between CG and CL (CG - CL, e) in butterfly species richness due to land abandonment for Red List species.

(a)

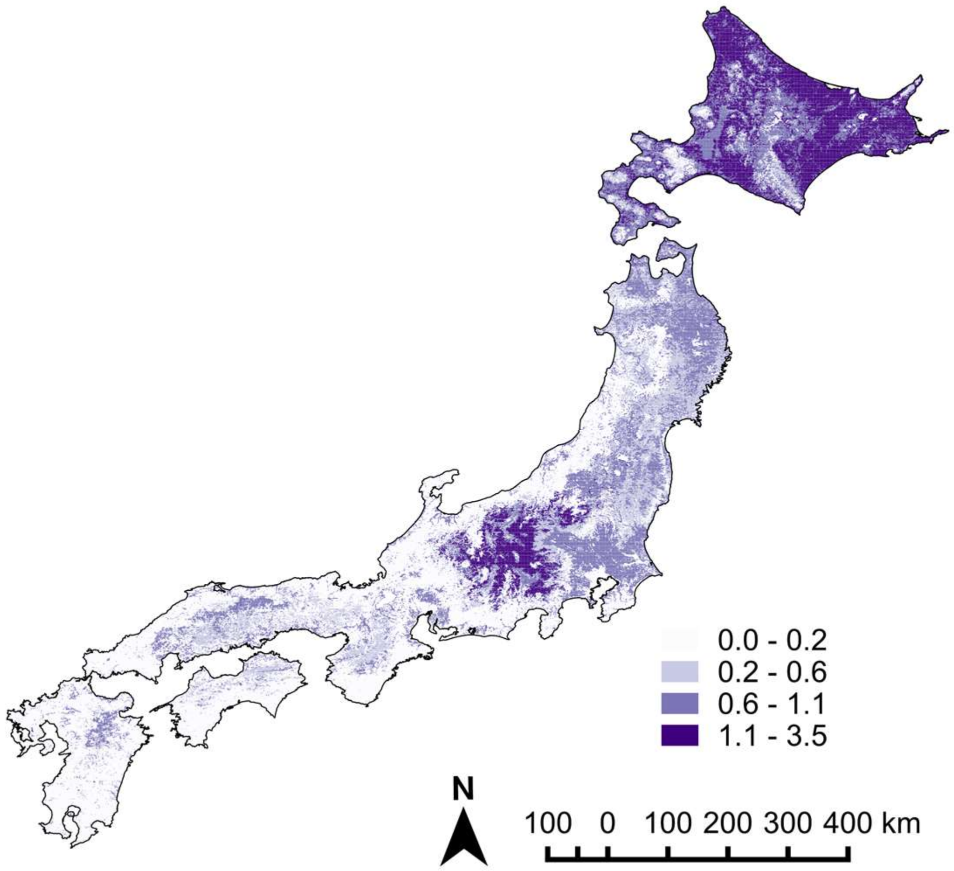


(b)

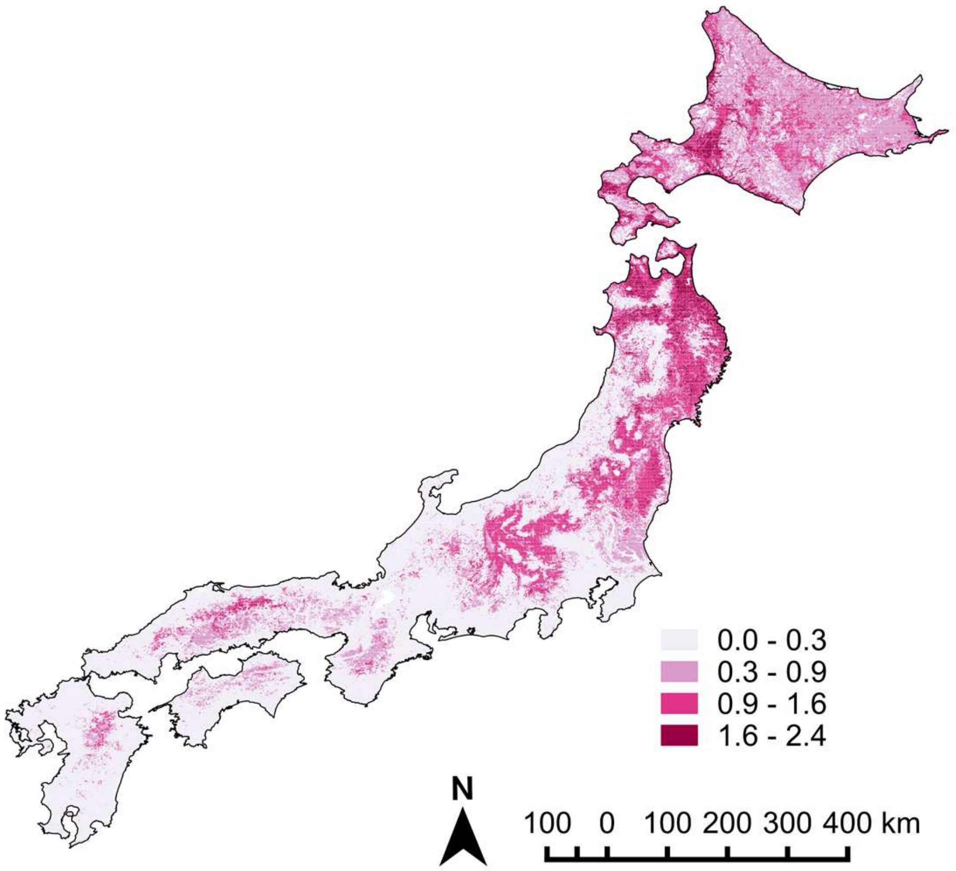

Fig. S4 continued.

(c)

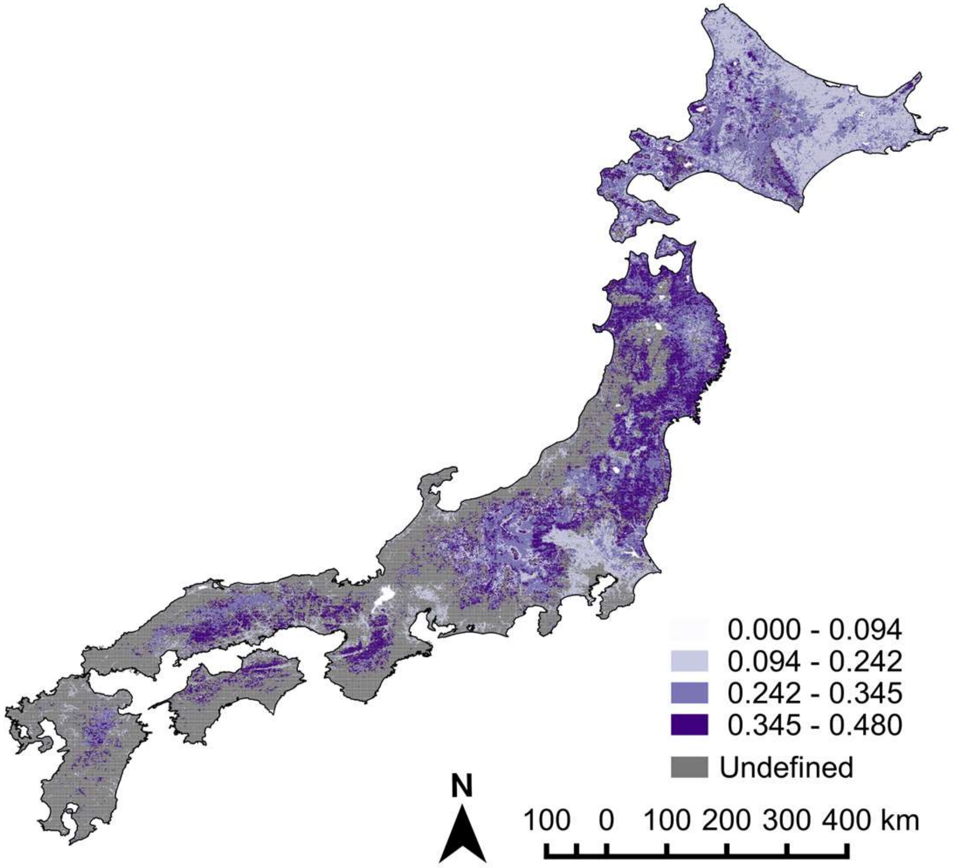


(d)

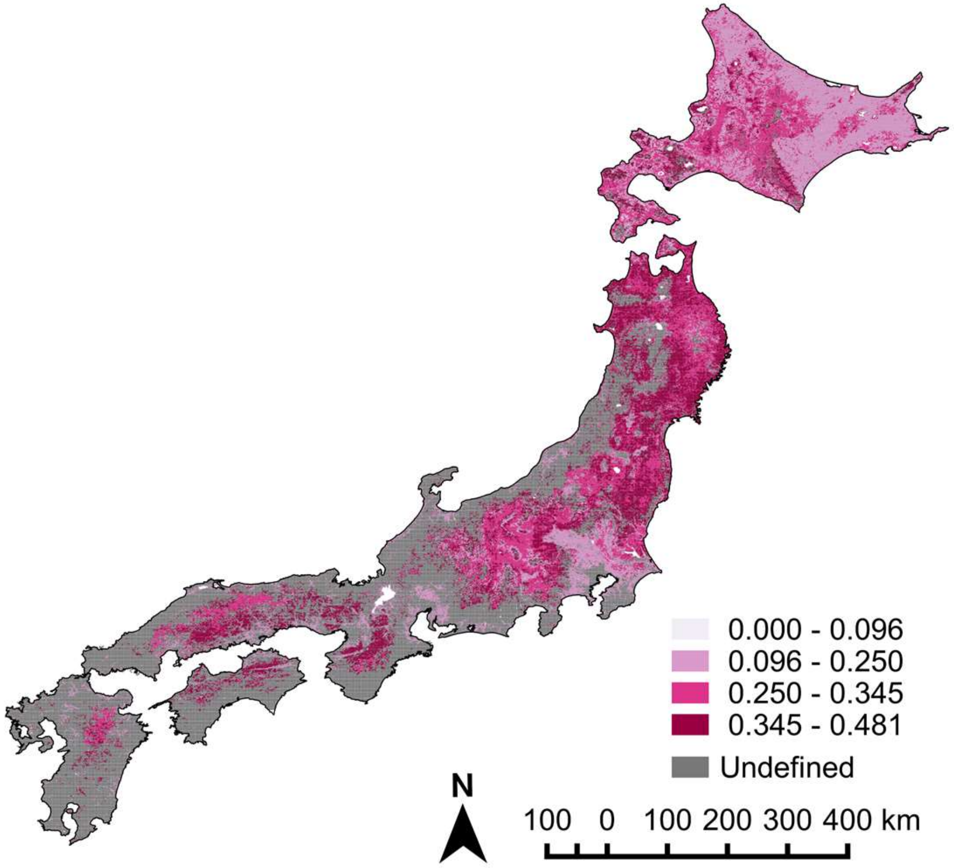

Fig. S4 continued.

(e)

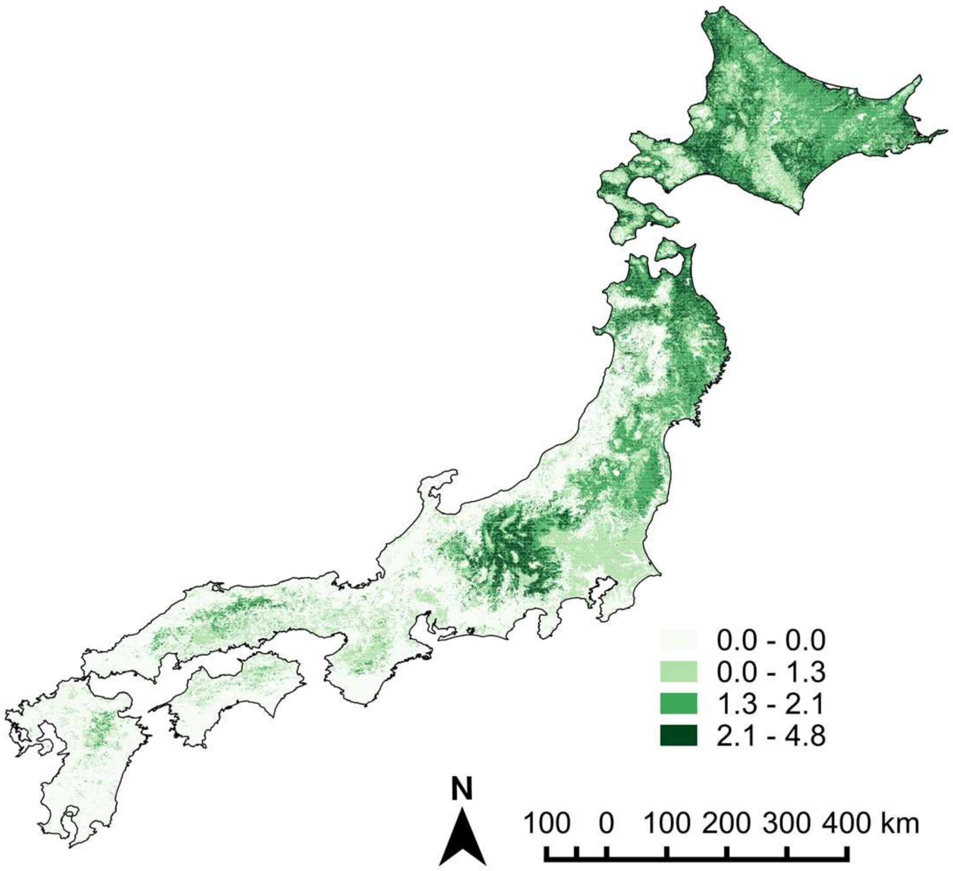
